# Root growth in Arabidopsis depends on the amount of glutathione and not the glutathione redox potential

**DOI:** 10.1101/2025.09.18.677034

**Authors:** Sajid A.K. Bangash, M. Taheb Safi, José M. Ugalde, Stephan Wagner, Kerstin A. Nagel, Marcus Jansen, Stephan Krueger, Markus Schwarzländer, Anna Moseler, Jean-Philippe Reichheld, Stanislav Kopriva, Andreas J. Meyer

## Abstract

Cell cycle activity and hence plant growth strictly depends on glutathione homeostasis. Despite compelling evidence for this dependency based on glutathione depletion, the cause for the cell cycle arrest had remained unclear. Cell cycle control may depend on either the absolute amount of glutathione or on the glutathione redox potential (*E*_GSH_). To unambiguously distinguish those two options, we characterized an allelic series of six Arabidopsis mutants affected in glutamate-cysteine ligase, which catalyses the first step for the biosynthesis of reduced glutathione (GSH). When grown under the same conditions, even mutants with 20% and 50% of wild-type GSH amounts are slightly stunted. The most severely compromised mutants, *zir1* and *rml1*, were crossed with either *gr1*, which lacks cyto-nuclear glutathione disulfide reductase and was used to induce a pronounced shift in *E*_GSH_, or with *bir6*, which has a diminished glutathione consumption and thus exhibits slightly increased levels of GSH. Based on theoretical considerations, these levels are not expected to shift the *E*_GSH_ to any significant extent. Our study shows that deleting GR1 in the *zir1* or *rml1* background does not result in an obvious phenotypic change. By contrast, deleting BIR6 was sufficient to suppress the growth arrest in *rml1* and to attenuate the growth in *zir1*. These findings demonstrate that root growth is dependent on the availability of sufficient amounts of GSH, and not affected by pronounced changes in *E*_GSH_. This insight provides a decisive step towards understanding the mechanisms underpinning the proposed role of glutathione in cell cycle and growth control.

**Highlight:** Crosses of GSH-deficient Arabidopsis mutants with mutants affected in glutathione reduction or GSH consumption pinpoint the GSH amount rather than the glutathione redox potential as causative for changes in root growth.

## Introduction

Plants have evolved sophisticated defence systems that allow them to withstand frequent adverse conditions with multiple biotic and abiotic stresses. Among the metabolic defence systems, the tripeptide glutathione (γ-Glu-Cys-Gly) plays a special role, fulfilling multiple functions in plant metabolism and stress defence. Several detoxification pathways for the removal of toxic metabolites, toxic xenobiotics and heavy metals rely on the reduced form of glutathione (GSH). These detoxification processes involve the conjugation of electrophilic metabolites with GSH for sequestration to the vacuole, formation of phytochelatins for binding and vacuolar sequestration of heavy metals, and binding of NO for subsequent reduction (Cassier-Chauvat *et al*., 2023; Noctor *et al*., 2011). GSH is an essential cofactor in the detoxification of formaldehyde and methylglyoxal. However, it is important to note that there is no net consumption of GSH in the respective reactions (Achkor *et al*., 2003; Rabbani *et al*., 2020). Similarly, GSH is essential as a cofactor for the coordination and protection of [2Fe-2S] clusters by glutaredoxins (GRXs) (Kumar *et al*., 2011; Moseler *et al*., 2015; Rouhier *et al*., 2007). A widespread GSH deficiency would obviously lead to severe impairment of all these processes and likely have pleiotropic consequences.

Glutathione constitutes the most abundant low molecular weight thiol and is present in cells at low millimolar concentrations (Meyer *et al*., 2001). While the two chemical species constituting the glutathione redox couple, i.e., the reduced form GSH and the oxidised form glutathione disulfide (GSSG), can readily be interconverted by oxidation and reduction, the major proportion of glutathione in unstressed cells is present as GSH, with only nanomolar concentrations of GSSG present in the cytosol, chloroplasts, mitochondria and peroxisomes (Meyer *et al*., 2007; Schwarzländer *et al*., 2008). In these four compartments, the extremely high level of reduction is maintained by glutathione disulfide reductases (GRs), and sufficient electron supply from NADPH to ensure continuous reduction of GSSG (Marty *et al*., 2019; Marty *et al*., 2009; Scherschel *et al*., 2024). The biosynthesis of GSH occurs through two enzymatic steps catalysed by glutamate-cysteine ligase (GSH1) and glutathione synthase (GSH2). Feedback inhibition occurs when the end product GSH acts on GSH1 (Hicks *et al*., 2007; Pasternak *et al*., 2008; Wachter *et al*., 2005; Yang *et al*., 2019). GSH1 has been shown to be the rate-limiting enzyme and several forward genetic screens have led to the isolation of weak recessive *gsh1* alleles (Ball *et al*., 2004; Cobbett *et al*., 1998; Jobe *et al*., 2012; Parisy *et al*., 2007; Shanmugam *et al*., 2012). Although the *cad2* (*cadmium-sensitive 2-1*), *pad2* (*phytoalexin-deficient 2*)*, rax1* (*regulator of APX2 1-1*) and *nrc1* (*non-response to cadmium 1*) mutants are all characterized by GSH levels of around 15-50% compared to the wild type (WT), stress-sensitive phenotypes only become apparent when they are exposed to cadmium, high light or pathogen stress. In contrast to these mutants, *zir1* (*zinc tolerance induced by iron 1*), containing about 15% of WT glutathione, is a semi-dwarf mutant even under control conditions, while remaining fully fertile. The apparent link between GSH amount and plant growth is in line with the far more severe *rml1* (*root meristemless 1*) allele, which contains less than 5% of WT GSH and shows strongly compromised postembryonic meristem activity in the primary root (Cheng *et al*., 1995; Vernoux *et al*., 2000). Null mutants devoid of endogenous GSH biosynthesis reach late stages of embryogenesis supported by maternal supply with GSH, but mutant seeds fail to germinate (Cairns *et al*., 2006; Lim *et al*., 2011). Detailed characterisation of the *rml1* mutant showed that the lack of GSH prevents progression through the cell cycle by blocking the G1-S transition. This effect can be phenocopied by treatment with *L*-buthionine-(S,R)-sulfoximine (BSO), a transition state analogue that blocks the active site of GSH1 and in turn inhibits GSH biosynthesis with high efficiency. A role for GSH in cell cycle control was recently demonstrated also in injured Arabidopsis roots, where GSH was recruited to cells near an ablation injury (Lee *et al*., 2025). Accumulation of GSH in these cells has been suggested to truncate the G1 phase, leading to coordinated entry into S phase (Lee *et al*., 2025). Depletion of GSH through incubation of Arabidopsis seedlings with BSO for 24 hours also caused a decrease in root hair length (Sánchez-Fernández *et al*., 1997). However, despite numerous efforts to elucidate the role of GSH in the cell cycle and root hair growth, the mechanistic involvement of GSH in these processes remains unclear.

The pronounced growth inhibition by BSO-induced GSH depletion was exploited to screen for BSO- resistant mutants. Such forward genetic screens identified the CLT transporters for GSH and its precursor γ-glutamylcysteine on the chloroplast envelope, which also mediate access of BSO to its nominal target GSH1 in the stroma, and BIR6, which was identified as a pentatricopeptide repeat protein responsible for splicing the mitochondrial genome-encoded transcript for NAD7, a subunit of complex I of the respiratory electron transport chain (Koprivova *et al*., 2010a; Maughan *et al*., 2010). Although a mechanism by which the lack of BIR6 ultimately causes partial resistance to BSO is still missing, the available data suggest that the *bir6* mutants maintain higher GSH levels on BSO at the whole-seedling level. It is unknown though, whether diminished consumption of GSH would also lead to increased steady state levels of GSH in the root meristem.

Because two molecules of GSH after oxidation form one molecule of GSSG, the glutathione redox potential (*E*_GSH_) is not only dependent on the GSH:GSSG ratio of the local glutathione pool but also on the total amount of glutathione. Therefore, GSH-deficient mutants are characterized by a distinct shift of their *E*_GSH_ towards less negative (i.e., less reducing) values. For the Arabidopsis mutant *rml1*, cytosolic *E*_GSH_ has been reported at about -260 mV compared with -315 mV in the cytosol of WT plants (Aller *et al*., 2013; Meyer *et al*., 2007; Schnaubelt *et al*., 2015). Schnaubelt *et al*. (2015) concluded that the altered redox potentials in the nucleus and cytosol of glutathione-deficient seedlings cause the cell cycle to arrest in roots, but not in shoots. A shift in *E*_GSH_ by more than 50 mV may be assumed to have a pronounced effect on class I glutaredoxins (GRXs) and their respective target proteins (Schlößer *et al*., 2024). GRXs can act as bidirectional redox operators, which mediate reversible post-translational modification of their respective target proteins, as shown for the redox-sensitive thiol/disulfide switch on the redox-sensitive green fluorescent protein 2 (roGFP2) (Geissel *et al*., 2024; Lang *et al*., 2024). Such post-translational thiol modification may be important for modulation of protein function (Mittler *et al*., 2022). In the moss *Physcomitrium patens*, deletion of the only chloroplast class I GRX causes a delayed recovery of the protein cysteinyl redox state after oxidative challenge (Bohle *et al*., 2024). Although specific target proteins of class I GRXs in plants remain unknown, similar functions in control of the protein redox state in the cyto-nuclear compartment can be assumed. In bacteria, class I GRXs have been shown to mediate electron transport from GSH to ribonucleoside-diphosphate reductase (RNR) as a reductive enzyme that converts ribonucleoside diphosphates to deoxyribonucleosides, which is essential for DNA synthesis and repair (Holmgren, 1976). Although it is not known whether plant RNRs function in a similar way to their bacterial counterparts, the process illustrates how severe aberrations in *E*_GSH_, in conjunction with class I GRXs, can affect key metabolic processes that are essential for maintaining normal cell cycle activity.

Considering the multifaceted roles of GSH in the cell, it has remained unresolved whether the effect of GSH on the cell cycle is predominantly dependent on *E*_GSH_ or on the absolute level of GSH. Such a distinction is desirable, however, since it would provide a criterion to focus on plausible candidate mechanisms that link glutathione and associated phenotypes. In order to resolve this question, we first characterised a comprehensive series of *gsh1* mutants alongside each other focussing on their root and shoot growth. We then developed a combinatorial genetic strategy that allowed us to titrate the absolute level of GSH with negligible effects on the redox state or, conversely, to shift the GSH:GSSG ratio in low-glutathione mutants without changing the glutathione amount and thus disentangle effects of *E*_GSH_ from effects mediated by the glutathione amount.

## Materials and methods

### Plant material and growth conditions

*Arabidopsis thaliana* (L.) Heynh. ecotype Columbia-0 (Col-0) was used throughout all experiments as WT. Mutants with defects in *GSH1* (*At4g23100*) were provided by the colleagues who reported their isolation and initial characterization (*cad2*, *rml1*: Chris Cobbett, *rax1*: Phil Mullineaux, *pad2*: Felix Mauch, *nrc1*: Julian Schroeder, *zir1*: Kuo-Chen Yeh). The null mutants *gr1* (Marty *et al*., 2009) and *bir6- 1* (Koprivova *et al*., 2010a) have been described earlier. Arabidopsis plants were grown on a mixture of soil (Floradur multiplication substrate, Floragard, Oldenburg, Germany), sand and perlite in 10:1:1 ratio and kept in controlled growth chambers under long day conditions with 16 h light at 19 °C and 8 h dark at 17 °C. Light intensity was kept between 50 and 75 µmol photons m^-2^ s^-1^ using fluorescent Philips TL-D 36W/865 Cool Daylight tubes and relative humidity at 50%. For phenotypic and physiological characterization, seedlings were grown on vertically oriented agar plates and sterile conditions. Seeds were surface-sterilized with 70% (v/v) ethanol for 5 min and plated on ½ MS medium (M0221; Duchefa Biochemie, Haarlem, The Netherlands), supplemented with 0.5% (w/v) sucrose and solidified with 0.8% (w/v) agar adjusted to pH 5.7–5.8 with KOH. Seeds were stratified for 2 d in the dark at 4 °C and germinated under long day conditions with 16 h light at 22 °C and 8 h dark at 18 °C. Light intensity was 75 µmol photons m^-2^ s^-1^ using fluorescent Osram L 18W/840 Cool White tubes and relative humidity at 50%.

### Genotyping

The *Arabidopsis thaliana* mutants *gr1-1* (*At3g24170*, SALK_105794, Nottingham Arabidopsis Seed Centre, Nottingham, UK), *bir6-1* (*At3g48250*), *zir1* and *rml1* (both *At4g23100*) were all described previously (Cheng *et al*., 1995; Koprivova *et al*., 2010a; Marty *et al*., 2009; Shanmugam *et al*., 2012). Homozygous *gr1-1* plants were identified by PCR using primers for the genomic locus (cs330, cs331) or for the insertion (LBb1.3, cs331) (Supplemental Table S1). Plants homozygous for *zir1*, *rml1* and *bir6-1* were screened by PCR with subsequent restriction analysis according to details provided in supplemental information (Supplemental Table S1). For crosses, double homozygous plants were selected in the F2 generation by PCR and restriction analysis.

### Phenotypic characterization

For determination of the root length, seedlings were documented on a stereomicroscope (Leica M165 FC, Leica Microsystems, Wetzlar, Germany) with a DFC425 camera roots analysed in Fiji (Schindelin *et al*., 2012). Further automatized phenotyping experiments were carried out using the automated platform GrowScreen-Agar at institute IBG-2: Plant Sciences, Forschungszentrum Jülich GmbH as previously described (Nagel *et al*., 2020). Root phenotyping was carried out using plates with solidified agar while shoot phenotyping used soil-grown plants. Arabidopsis seeds were surface-sterilized with sodium hypochlorite and then sown on one-third Hoagland’s solution, solidified with sterile agarose in Petri dishes (120 × 120 × 17 mm) (Nagel *et al*., 2020). Seeds were stratified at 4 °C in the dark for 5 d before they were grown vertically in a climate chamber set to long-day conditions as previously described (Nagel *et al*., 2020). Petri dishes were placed in the automated platform GrowScreen-Agar and images of the root systems were taken automatically via a high-resolution CCD-camera (IPX-6 M3- TVM, Imperx Inc., Boca Raton, FL, USA) under infrared illumination (Caliandro *et al*., 2013). Root system architecture of each plant was then analysed by the GROWSCREEN-ROOT method (Nagel *et al*., 2009).

Shoot growth was analysed automatically by using the GROWSCREEN FLUORO setup described earlier (Barboza-Barquero *et al*., 2015; Jansen *et al*., 2009). Phenotyping of *gsh1* mutants was performed as previously reported (Bangash *et al*., 2019).

### In vivo visualization of the glutathione redox potential with roGFP2

To enable in vivo determination of relative changes in the glutathione redox potential WT plants and all *gsh1* mutants were transformed with Grx1-roGFP2 targeted to the cytosol. For this, Grx1-roGFP2 was cloned behind the Arabidopsis *ubiquitin-10* promoter in the binary vector pBinCM, which was constructed from the pBin19 derivative pBinAR (Aller *et al*., 2013). Reporter lines with ubiquitous expression were selected through a manual optical screen on a fluorescent stereomicroscope (M165 FC, Leica Microsystems, Wetzlar, Germany). Stable lines homozygous for the reporter construct were generated through two rounds of self-fertilization and selection of lines with close to 100% fluorescent progeny. Ratiometric imaging of Grx1-roGFP2 was done on 5-day-old seedlings. Seedlings were mounted on a slide in a drop of water, covered by a cover slip and immediately transferred to the stage of a confocal microscope (LSM780, Carl Zeiss Microscopy, Jena, Germany). Images of root tips were collected with a 40× lens (Zeiss C-Apochromat 40×/1.2 NA water immersion). roGFP2 was consecutively excited at 405 and 488 nm, respectively, in multi-track mode with line switching. All lines were scanned four times and the signal averaged to improve the signal-to-noise ratio in the final images. Emission was collected between 504 and 530 nm. Autofluorescence excited at 405 nm was collected from 431 to 470 nm. The relative power output for the laser lines 405 nm and 488 nm was always kept at 1:3 ratio. Images were processed using a custom MATLAB tool (Fricker, 2016) with x,y noise filtering, fluorescence background subtraction and autofluorescence correction as described previously (Lehmann *et al*., 2009).

### In vivo visualization of glutathione with monochlorobimane

Visualization of glutathione in root tips of 5-day-old seedlings was performed as previously described (Meyer *et al*., 2001). Seedlings were incubated in 100 µM monochlorobimane (MCB, Molecular Probes, Thermo Fisher Scientific) in deionized water for 30 min, followed by 5 min of incubation in 50 µM propidium iodide (PI, Thermo Fisher Scientific). Seedlings were then washed in deionized water and mounted on a slide for image acquisition. Z-stacks of root tips were collected with a 25× lens (Zeiss LD LCI Plan-Apochromat 25x/0.8 NA water immersion). Fluorescence of GSB and PI were simultaneously excited at 405 nm and 543 nm, respectively. Emission of GSB was collected for the band 449-613 nm and emission of PI between 613 nm and 704 nm, respectively. Maximum intensity projections of z-stack images were generated in Fiji (Schindelin *et al*., 2012) and exported as merged and individual channel images. To quantify bimane fluorescence, the relevant fluorescence channel of the maximum projection was segmented for the root and the fluorescence intensity in this area was measured in Fiji.

### Analysis of low-molecular weight thiols

20 mg plant material was homogenised and extracted in a 10-fold volume of 0.1 N HCl. Samples were centrifuged for 10 min at 14,000 *g* and 4 °C. 25 µL of the supernatant were mixed with 20 µL of 0.1 M NaOH and 1 µL of 100 mM freshly prepared dithiothreitol to quantitatively reduce disulfides. Samples were vortexed, spun down and kept for 15 min at 37 °C in the dark. Afterwards, 10 μL 1 M Tris/HCl pH 8.0, 35 μL water and 5 μL 100 mM monobromobimane in acetonitrile (Thiolyte® MB, Calbiochem, Merck Millipore) were mixed and added to the samples. The samples were vortexed, spun down and kept for 15 min at 37 °C in dark before derivatization was stopped by adding 100 μL of 9% (v/v) acetic acid. Subsequently, samples were vortexed and centrifuged at 13,000 *g* for 15 min at 4 °C. 180 µL of the supernatant were filled in HPLC vials. Thiol conjugates were separated by HPLC (SpherisorbTM ODS2, 250 × 4.6 mm, 5 µm, Waters) using buffer C (10% (v/v) methanol, 0.25% (v/v) acetic acid, pH 3.7) and D (90% (v/v) methanol, 0.25% (v/v) acetic acid, pH 3.9). The elution protocol was employed with a linear gradient from 4 to 20% D in C within 20 min, with the flow rate set to 1 mL/min. Bimane adducts were detected fluorimetrically (474 detector, Waters) with excitation at 390 nm and emission at 480 nm.

### Extraction and quantification of amino acids

Eight-day-old seedlings were used for amino acids quantification. Amino acids were extracted from 100 mg frozen plant material in 400 µL of 80% (v/v) ethanol by incubation for 15 min at 4 °C on a thermomixer operated at 600 rpm. After incubation, extracts were centrifuged for 10 min at 4 °C and 21,000 *g*. The supernatant was transferred to fresh tubes and stored on ice. The remaining pellet was extracted again with 400 µL of 80% (v/v) ethanol, repeating the previous step. The pellet was re-extracted for a third time and supernatants of all three extractions were combined and either directly used for HPLC measurements or stored at -20 ℃ until further use.

For amino acids analysis, ethanolic extracts were diluted 1:10 with water before loading into the HPLC autosampler (Ultimate Autosampler 3000; Dionex, Thermo Fischer). Prior to separation, the amino acids were derivatised in a separate reaction vial by pre-column derivatisation consisting of an automated mixing of 25 µL of 1× o-phthalaldehyde solution (Grace Davison Discovery Sciences, Carnforth, UK), 5.5 µL of 1M borate buffer pH 10.7 and 25 µL of sample immediately prior to analysis. After 90 s incubation, 20 µL of the derivatisation mixture were injected into the HPLC system and separated on a HyperClone 3 µm ODS(C18)120 150×4.6 mm HPLC column (Phenomenex) by a discontinuous gradient consisting of solvent A (8.8 mM sodium phosphate (NaPO_4_) pH 7.5 and 0.2% (v/v) tetrahydrofuran) and increasing proportions of solvent B (18.7 mM NaPO_4_ pH 7.5, 32.7% (v/v) methanol (MeOH) and 20.6% (v/v) acetonitrile). Separated fluorescent derivatives were detected at 480 nm after excitation by 380 nm using a RF 2000 fluorescence detector (Dionex) in the "low" sensitivity mode. Data acquisition and analysis was done using the Chromeleon software (6.80 SR7; Dionex). For peak identification and quantification, various proteinogenic amino acids (Sigma-Aldrich) were used as standards.

### GSH1 protein homology modelling

A homology model of Arabidopsis GSH1 was built by using MODELLER (http://salilab.org/modeller/) with *Brassica juncea* GSH1 (BjGSH1; PDB: 2GWC; 96% sequence identity) as a template. The transition state analogue and ADP molecules shown in the structure were copied from *E. coli* GCL (PDB: 2D33) after superimposition with the *At*GSH1 model.

## Results

### GSH-deficient mutants display growth phenotypes

Mutants that are partially glutathione-deficient, with intermediate GSH levels ranging from 15% to 50% of wild-type glutathione levels, were described as wild-type-like in appearance by several studies (Ball *et al*., 2004; Howden *et al*., 1995; Parisy *et al*., 2007). However, a side-by-side comparison of the *gsh1* mutants *cad2*, *pad2* and *rax1* with wild-type controls indicated that all three mutants had smaller shoots when grown under the same conditions on agar plates for 14 days (Schnaubelt *et al*., 2015). When grown in soil for six weeks, *pad2* and *rax1*, but not *cad2*, were smaller than the WT (Bangash *et al*., 2019). In order to elucidate the role of glutathione in plant growth, here with a particular focus on root growth, we assembled the full set of six known *gsh1* mutants. All of these mutants carry point mutations, except c*ad2*, which carries a six-base-pair out-of-frame deletion resulting in the loss of two amino acids and a V-to-L substitution in the subsequent position (**Fig. 1A**). Mapping these mutations onto the GSH1 structure that was resolved based on the closely related *Brassica juncea* GSH1 (Hothorn *et al*., 2006), shows that all the mutations surround the active site of the protein with *zir1* and *rml1* in close proximity to the substrate entry side (**Fig. 1B**). Side-by side comparison of all available *gsh1* mutants for their seedling development during the first five days after stratification, revealed only minor differences in root length between Col-0, *rax1*, *pad2*, *cad2* and *nrc1*, but significantly shorter roots for *zir1* and complete abolition of post-germination root growth in *rml1* (**Fig. 1C, D**). At 20 days after stratification, the total length of the roots, i.e., the sum of primary and all lateral roots, of *rax1*, *pad2*, *cad2* and *nrc1* was shorter than Col-0 roots. However, this decrease was largely caused by a decrease in the length of lateral roots and a trend towards a lower number of lateral roots (**Fig. 2A-C**). The roots of *zir1* were significantly smaller than all other mutants in all parameters. Because the growth of *rml1* seedlings stops immediately after germination, they were not analysed here. Similar phenotypic differences were also observed for shoots from soil-grown plants, which did not show a difference in the projected leaf area (PLA) seven days after stratification (**Fig. 2D**). However, after 20 days, *rax1*, *pad2*, *cad2* and *nrc1* were smaller than Col-0. While those four mutants were indistinguishable in size between each other, *zir1* was smaller at this point, reaching only about 30% of the wild-type PLA (**Fig. 2E**). Further morphological characterisation of shoot growth indicated a tendency for shorter petioles of GSH-deficient mutants, resulting in increased shoot compactness after 20 days (**Fig. 2F**).

**Fig 1.**
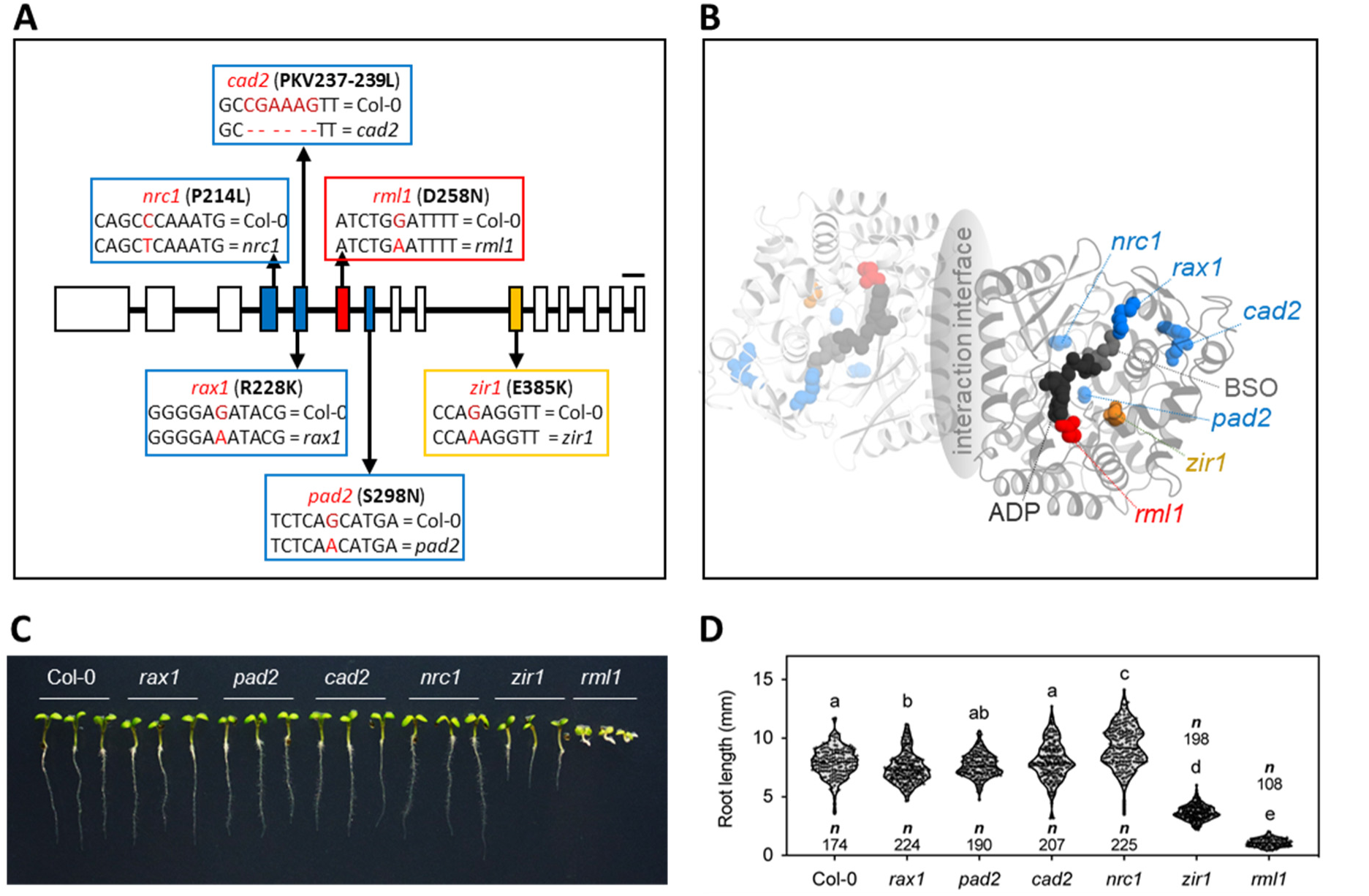
Allelic series of *gsh1* mutants in *Arabidopsis thaliana*. (A) Physical map of the Arabidopsis *GSH1* gene model (*At4G23100*) annotated for the known mutations. Exons and introns are illustrated as boxes and lines, respectively. Scale bar = 200 bp. The mutations at the DNA level are indicated in red letters and the resulting changes in the protein sequence is shown behind the mutant names. (B) Protein homology model of Arabidopsis *GSH1* depicting the physical position of all mutations relative to the active site (blue, orange and red balls). The model was built based on *Brassica juncea* GSH1 (PDB 2GWC). In addition to the mutations, the model also depicts BSO (*L*-buthionine-(S,R)-sulfoximine) (grey balls) as a transition state analogue, which inhibits GSH1 activity, and the cofactor ADP (black balls). (C) Phenotypes of 5-day-old seedlings of all *gsh1* mutants. The seeds were germinated and grown on ½ MS agar plates under a 16-hour light/8-hour dark cycle. (D) Root length of seedlings five days after germination. The letters represent statistical differences between genotypes according to a One-way ANOVA comparison, using a Tukey test (p<0.01).

**Fig. 2.**
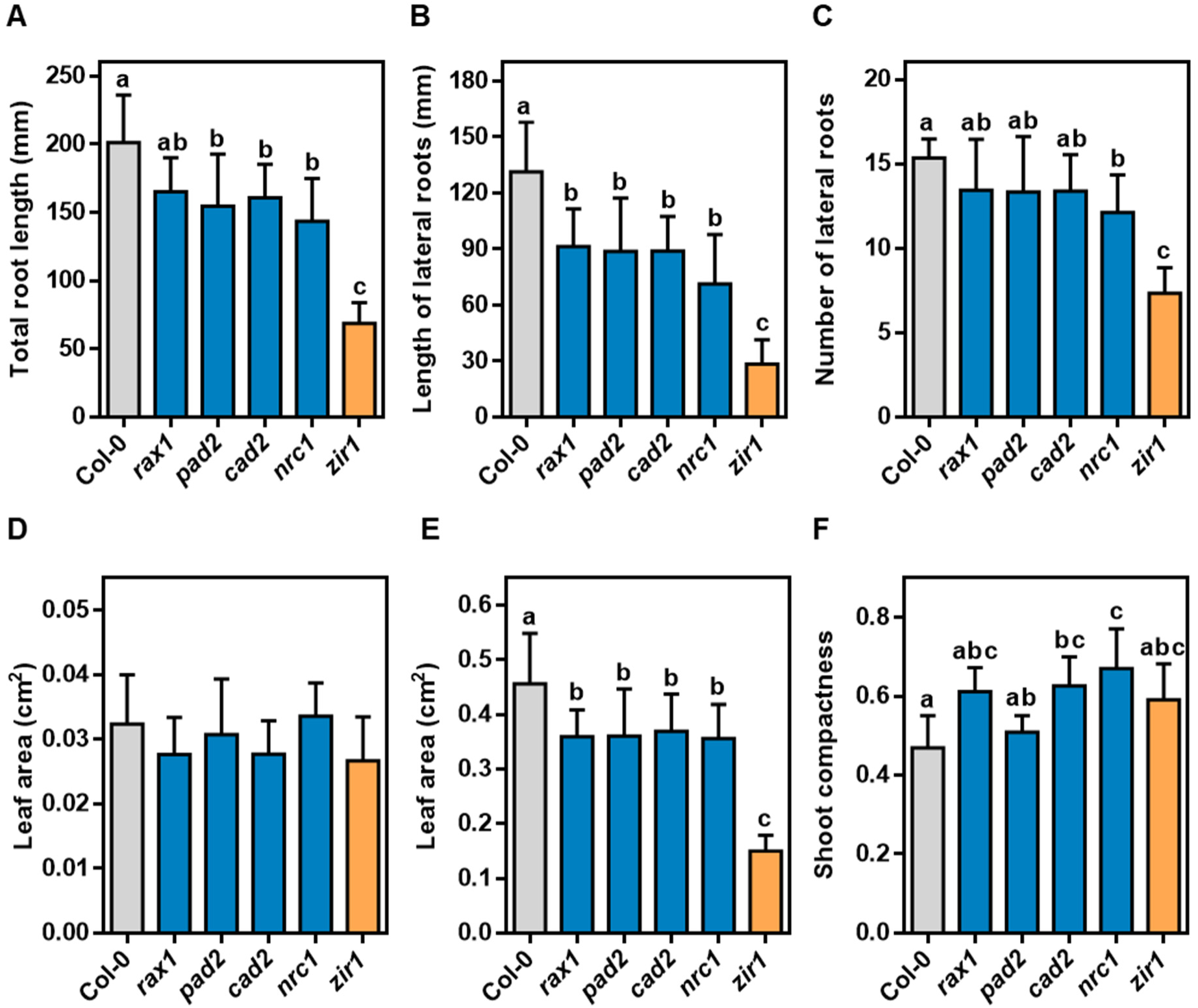
Root and shoot phenotypes of the allelic series of *gsh1* mutants. (A-C) Phenotypic characterization of roots grown in agar 20 days after stratification. The total root length, i.e., the sum of primary and all lateral roots (A), the length of lateral roots (B), and the number of lateral roots (C) were determined with the automatized root phenotyping platform GrowScreen-Agar. (D, E) Projected leaf area of soil-grown plants on day 7 (D) and day 20 (E) after stratification. (F) Shoot compactness 20 days after stratification. All values are means ± SD (n ≥ 10). The letters in each graph indicate significant differences (One-way ANOVA with Tukey’s multiple comparisons test; p < 0.05).

### Glutathione deficiency causes a shift in glutathione redox potential towards less reducing values

The general characteristics of glutathione deficiency is known for the entire allelic series of *gsh1* mutants. However, the glutathione levels reported in the literature are difficult to compare as individual mutants were grown under different conditions rather than analysed side by side. To compare the different mutants in a standardized manner side-by-side and to ultimately better understand the role of glutathione in determining growth, we analysed the entire allelic series for low molecular weight thiols and the glutathione redox potential (*E*_GSH_). We first labelled root tips with monochlorobimane (MCB) for GSH detection. Since glutathione disulfide reductase (GR) is active and GSSG is continuously being reduced to GSH, the formation of the fluorescent glutathione-bimane conjugate (GSB) provides close to quantitative labelling of the entire glutathione pool (Meyer *et al*., 2001). Compared to the WT, the GSB signal intensity was lowered in *rax1*, *pad2*, *cad2* and *nrc1* to fluorescence intensities of approximately 20-40% (**Fig. 3A, B**). When imaged with identical instrument settings, GSB in the root tips of *zir1* accumulated to about 15% compared to the WT, but detected fluorescence was close to background in *rml1*. Simultaneous staining with propidium iodide (PI) resulted in cell wall labelling only, with no significant nuclear staining, indicating that all cells in the root tips of all mutants were alive. The differences in GSB fluorescence in root tips were largely consistent with the amounts of GSH quantified by HPLC in whole seedlings (**Fig. 3C**). Conversely, the cysteine content, which is typically very low in WT seedlings, increased proportionally in all GSH-deficient lines (**Fig. 3D**). Yet, the *E*_GSH_ depends on both the absolute amount of glutathione and its redox state. The fluorescence ratio calculated for the biosensor Grx1-roGFP2, stably expressed in the cytosol of all mutant lines and the WT, first indicated that roGFP2 was almost completely reduced in the root tips of wild-type plants (**Fig. 3E**), which corresponds to an *E*_GSH_ of -315 mV or even more negative(Meyer *et al*., 2007). Compared to the WT, the fluorescence ratio of roGFP2 increased in the root tips of *rax1*, *pad2*, *cad2* and *nrc1*, while there was no difference between all these four mutants. The fluorescence ratio was much higher in *zir1* and especially in *rml1*, indicating a predominantly oxidised state of roGFP2. Given the accepted midpoint redox potential of -280 mV for roGFP2(Hanson *et al*., 2004), the intermediate fluorescence ratios coded in green with some red pixels interspersed allow to estimate an *E*_GSH_ in the order of about -275 mV. Based on the high fluorescence ratio approaching maximum oxidation in the cytosol of *rml1* root tip cells, one can estimate an *E*_GSH_ of approximately -260 mV, which is in good agreement with quantitative measurements in a previous report (Aller *et al*., 2013).

**Fig. 3.**
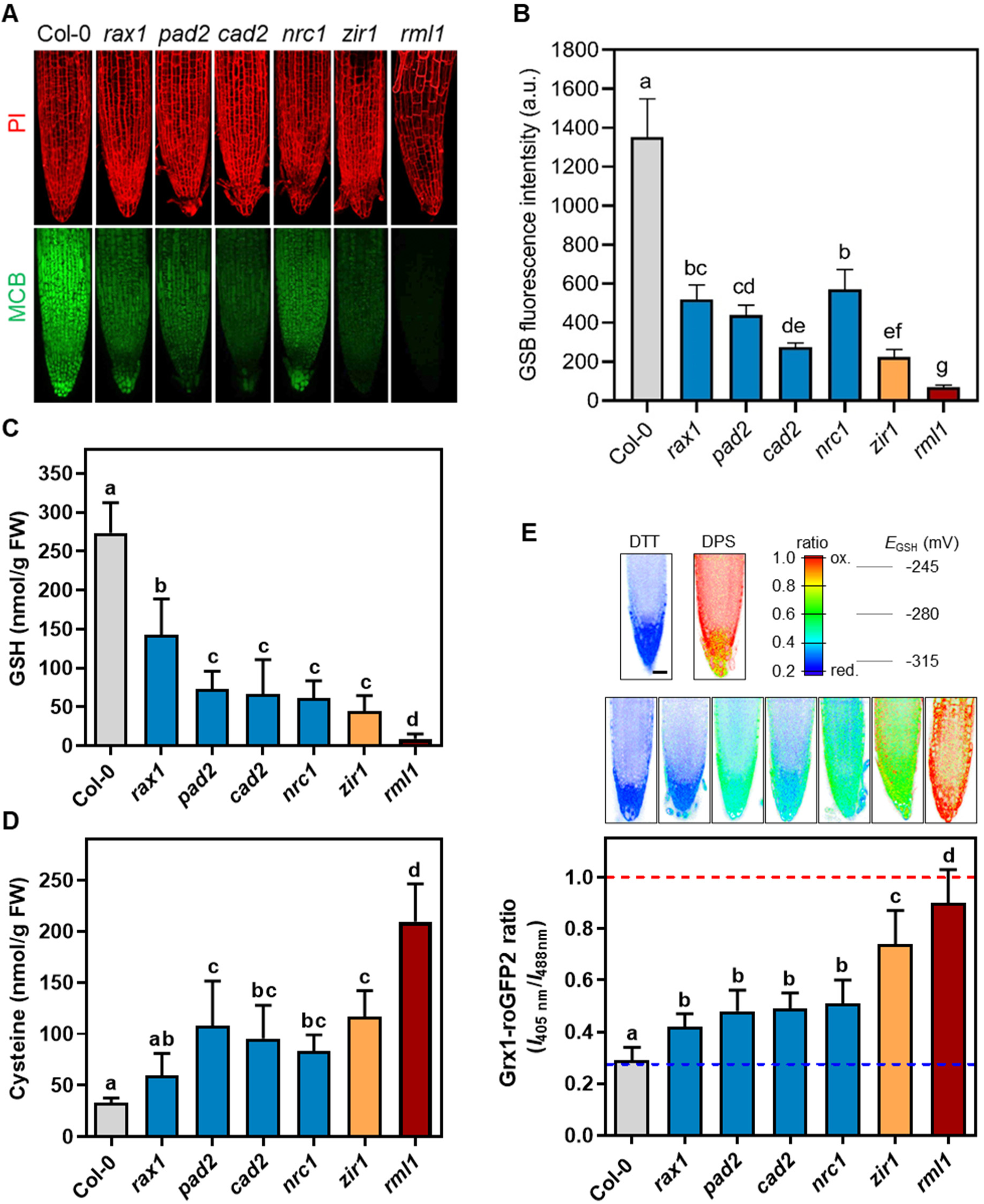
Quantitative analysis of low molecular weight thiols and the glutathione redox potential in *gsh1* mutants. (A) Fluorescent labelling of GSH in root tips. Five-day-old seedlings were labelled with propidium iodide (PI, 50 µM, red) to show plasma membrane integrity and monochlorobimane (MCB, 100 µM, green) for GSH conjugation. Stacks of images were collected by confocal laser scanning microscopy and are presented as maximum projections with the same instrument settings for all lines. The images show the fluorescence 30 min after start of the labelling. (B) Quantitative analysis of GSB fluorescence after labelling of root tips with 100 µM MCB for 30 min. All values are means ± SD (n ≥ 7). Letters in each graph indicate statistically significant differences (One-way ANOVA with Fisheŕs LSD test; p < 0.05). (C, D) HPLC analysis of glutathione (C) and cysteine (D). (E) Live cell imaging of the glutathione redox potential *E*_GSH_ with cytosolic Grx1-roGFP2 in root tips. Biosensor calibration was done with 10 mM dithiothreitol (DTT) for full reduction and 5 mM 2-2’-dipyridylsulfide (DPS) for full oxidation. To allow an estimation of *E*_GSH_ values from the ratio images, *E*_GSH_ values that limit the near linear range of the roGFP2 titration curve, i.e., ±35 mV around the known midpoint potential of -280 mV, are indicated (for details see (Meyer and Dick, 2010). For a semi-quantitative description of the redox state of roGFP2, projected images generated from an image stack were analysed for the 405 nm/488 nm excitation ratio. All values are means from ≥ 10 biological replicates; error bars = SD. Scale bar = 20 µm. Letters in each graph indicate significant differences (One-way ANOVA with Tukey’s multiple comparisons test; P < 0.05).

Partial suppression of the *rml1* root growth deficient phenotype can be achieved by external supply of 250 µM GSH (Vernoux *et al*., 2000). By titrating with different concentrations of GSH we found that 50 µM GSH was sufficient to reactivate meristem activity in some *rml1* roots, although the degree of the reactivation remained limited (**Fig. 4A, B**). External supply of 1 mM GSH resulted in a stronger reactivation of root growth and larger shoots of both *rml1* and *zir1* (**Fig. 4C, G**).

**Fig. 4.**
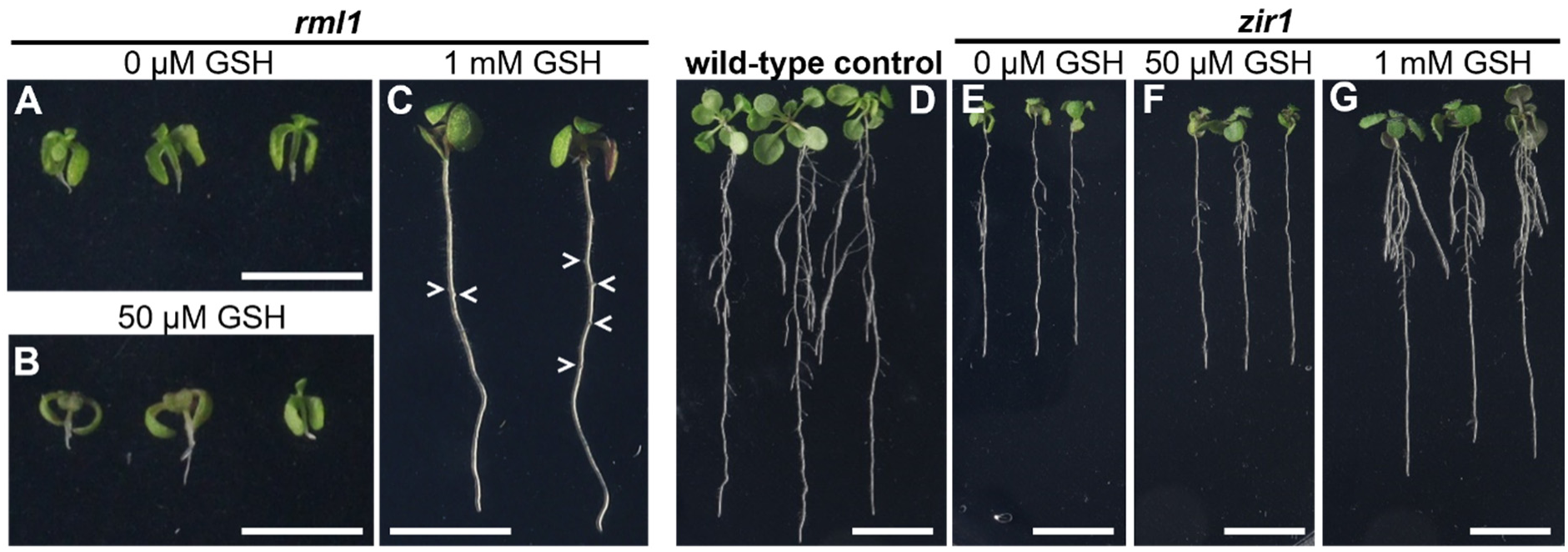
Feeding low amounts of GSH is sufficient to partially suppress root growth deficiency in *rml1* and *zir1*. (A-C) 12-day-old *rml1* seedlings grown either without GSH (A), with 50 µM GSH (B) or with 1 mM GSH (C). Arrowheads point at early lateral root primordia. (D) Wild-type seedlings grown without any supplemented glutathione. (E-G) 12-day-old *zir1* seedlings grown either without GSH (E), with 50 µM GSH (F) or with 1 mM GSH (G). After sterilisation, the seeds were stratified at 4 °C for two days and grown on ½ MS agar plates for four days. On day four, the seedlings were transferred to new plates containing the specified concentrations of GSH, where they were grown for a further eight days. Scale bars: *rml1* (panels A-C): 5 mm; WT and *zir1* (panels D-G): 10 mm.

### Severe GSH deficiency causes a global accumulation of free amino acids

Because free amino acids may become toxic at higher concentrations, amino acid homeostasis has to be regulated by controlling metabolic flux, which at the level of amino acid synthesis is mainly achieved by allosteric feedback inhibition (Hildebrandt, 2018). Given the pronounced increase of cysteine levels in *zir1* and *rml1* mutants (**Fig. 3C**), we investigated whether other amino acid levels were also increased to levels that might be considered toxic. While levels in *zir1* were similar to the wild-type control for most free amino acids, homozygous *rml1* seedlings showed a pronounced increase in all amino acids except for glutamine (**Fig. 5A**, Supplemental Table S2). This increase reached values of approximately 200% compared to the WT for most amino acids and was particularly evident when the relative abundance of the different amino acids was calculated (**Fig. 5B**). The accumulation of free amino acids may indicate that growth processes have largely ceased in *rml1*, resulting in reduced rates of protein synthesis and at the same time, increased proteolysis.

**Fig. 5.**
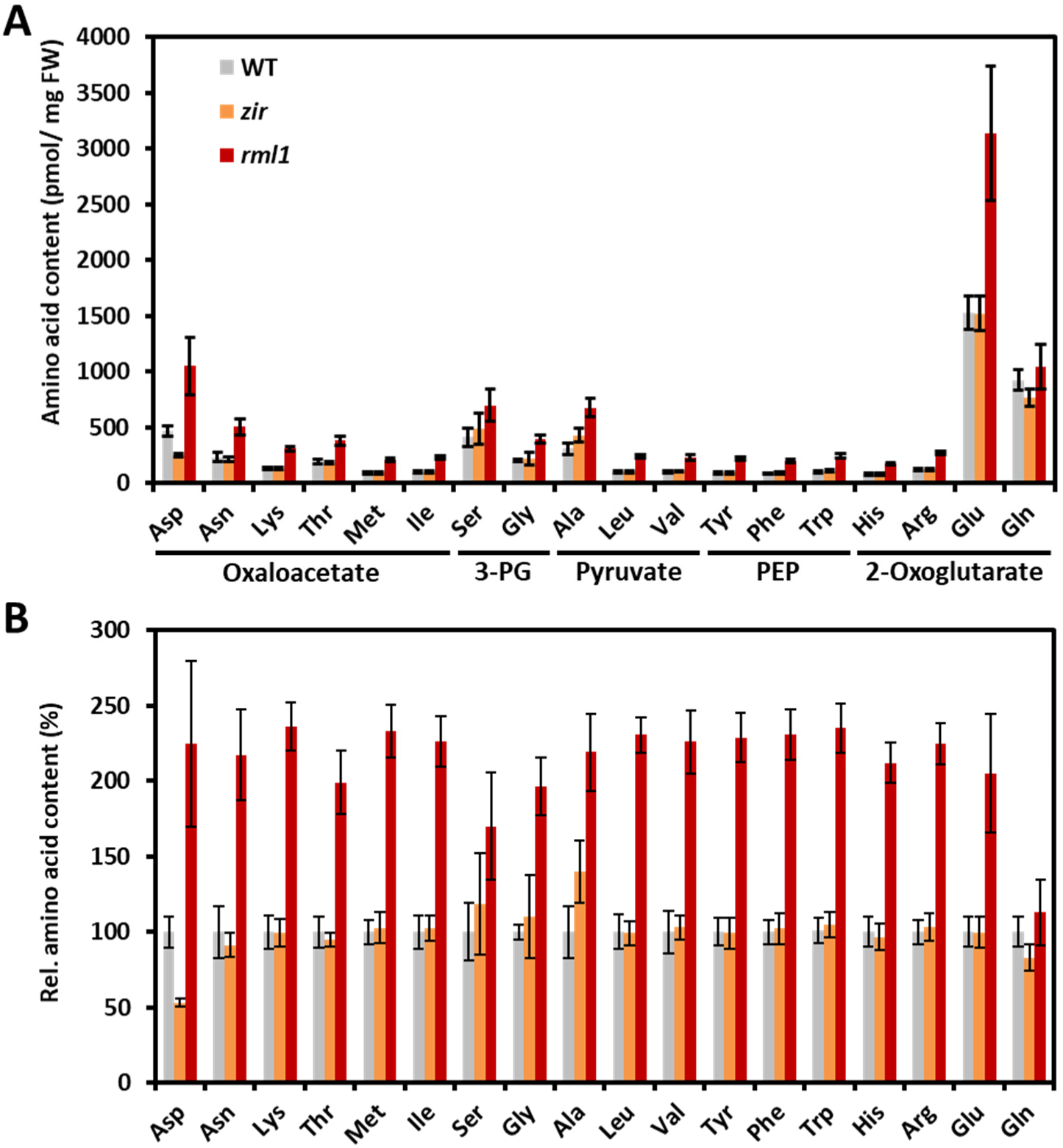
Severe GSH deficiency in *rml1* leads to accumulation of amino acids. (A) The amino acids were analysed in whole seedling extracts collected from seedlings grown for eight days under axenic conditions with a 16-hour light/8-hour dark illumination regime. The amino acids are grouped according to their common precursor. 3-PG: 3-phosphoglycerate; PEP: phosphoenolpyruvate. (B) Relative amounts of amino acids of *zir1* and *rml1* compared to the wild-type control. The values for the WT were set to 100%. Values are means ± SD (*n* = 5).

### The absolute amount of GSH present in the cyto-nuclear compartment defines root growth

Although all experiments strongly suggest that the glutathione availability is a key determinant of growth, the analyses do not allow for any unambiguous distinction between the absolute amount of glutathione and the resulting *E*_GSH_ as the decisive cause. To disentangle these two parameters, we designed a genetic assay in which we crossed GSH-deficient mutants with either a *gr1* mutant, which lacks cyto-nuclear GR activity and thus the ability to efficiently reduce GSSG to GSH (Marty *et al*., 2009), or *bir6-1*, a mutant with lower GSH turnover, which is anticipated to maintain a higher steady-state level of GSH as a consequence. Due to the 2:1 stoichiometry in the redox conversion of glutathione, it is important to note that *E*_GSH_, as calculated by the Nernst equation, is far more sensitive to changes in GSSG than to changes in GSH or the total glutathione pool. The rationale is that mutants with low GSH and lacking GR1 would exhibit an additive phenotypic effect if *E*_GSH_ were the causative factor. In the most extreme case, a lack of both sufficient GSH and GR1 activity could be fatal (**Fig. 7A**). Conversely, double mutants with severely impaired biosynthesis but decreased GSH consumption would exhibit an elevated steady-state concentration of GSH in the cytosol and the nucleus, provided that the steady-state concentration is set by the rates of GSH biosynthesis and consumption. If the steady-state level of GSH was critical, we expected to see suppression of the *gsh1* dwarf phenotype in such a combination of mutants (**Fig. 7A**).

To validate the *bir6* as a model for higher GSH levels than wild-type controls when grown on *L*-BSO, 3- day-old seedlings were transferred to plates supplemented with or without BSO and grown for a further four days. Under control conditions without BSO the primary roots of *bir6* seedlings were shorter than those of wild-type seedlings. However, when grown on 1 mM *L*-BSO, wild-type seedlings were clearly shorter than the respective roots grown without BSO. For *bir6* primary roots no such growth inhibition was observed after four days on BSO (**Fig. 6A**). This result shows that after four days on BSO growth inhibition caused by BSO was only apparent in wild-type seedlings but not in *bir6* seedlings. Consistently, the primary root tips of control seedlings grown without BSO did not exhibit a pronounced difference in fluorescence intensity after GSH labelling with MCB (**Fig. 6B** left panel). Yet, a clear difference in MCB labelling was observed in root tips after four days on BSO. Although the MCB labelling was largely diminished, the BSO-treated *bir6* roots showed residual GSB fluorescence, whereas GSB fluorescence was indistinguishable from background signal in BSO-treated wild-type root tips (**Fig. 6B** right panel). In all cases, the absence of PI labelling of nuclear DNA indicated the integrity of cell membranes and maintained cell viability.

**Fig. 6.**
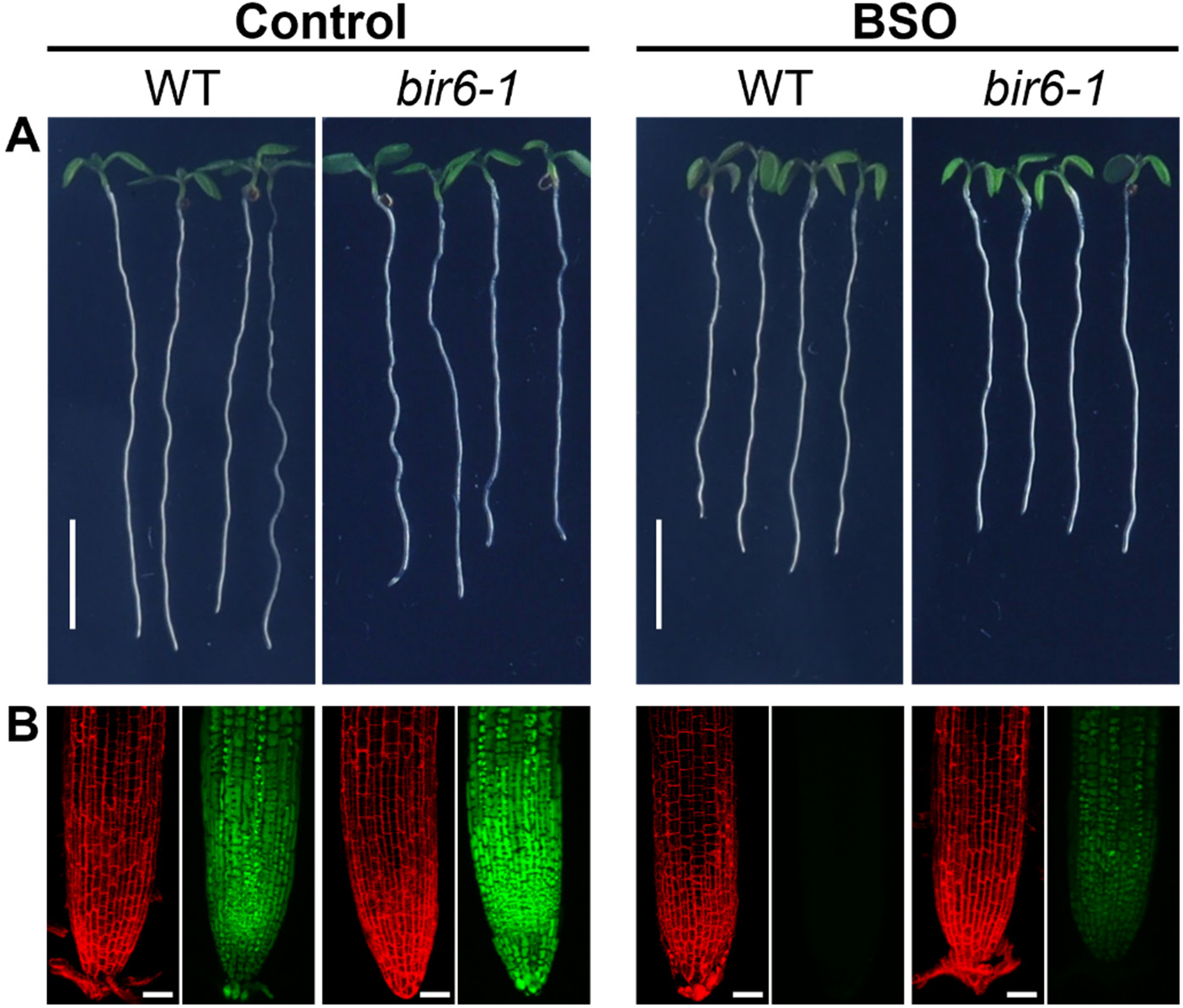
BSO-induced depletion of GSH in primary root tips of *bir6* is less severe than in wild-type roots. (A) Primary root length of seedlings grown on medium with or without BSO. After an initial stratification for two days at 4 °C, seedlings were grown on ½ MS agar plates for three days and then transferred to new plates without BSO (control) or supplemented with 1 mM BSO and grown for further four days. Scale bars = 5 mm. (B) Labelling of GSH in primary root tips. Seedlings were labelled with 100 µM monochlorobimane (MCB) for GSH (green) and 50 µM propidium iodide (PI) for cell walls (red) and cell membrane integrity (absence of DNA staining) for 30 min. The depicted images are maximum projections generated from stacks of images. For imaging of residual MCB fluorescence in BSO-treated roots the power of the 405 nm laser line was doubled compared to settings used for control seedlings. Scale bars = 20 µm.

Finally, we aimed to resolve whether root growth in severely GSH-deficient mutants is limited by a shift in *E*_GSH_ towards less negative (i.e., less reducing) redox potentials or by the total GSH amount in cells of the meristematic region in the root tip. To this end, we performed crosses of *rml1* and *zir1* with *bir6* and *gr1* (**Fig. 7A**). Double homozygous seedlings were selected by PCR (**Fig. 7B**). While *zir1* crosses were all viable and fertile in the double homozygous state, *rml1* mutants could not be maintained in the double homozygous form. Therefore, experiments had to be carried out on seedlings selected for their characteristic dwarf phenotype that segregated from seeds generated on a *bir6*^-/-^ *rml1*^+/-^ plant after self-fertilization. Comparison of dwarf seedlings segregating from a *bir6 rml1* population with homozygous *rml1* seedlings showed two striking, macroscopically visible phenotypes. First, the primary roots of *bir6 rml1*, although still stunted, were slightly longer than those of *rml1* (**Fig. 7C** top row). Second, the primary roots also grew root hairs, which are severely stunted or even absent in *rml1* (**Fig. 7C** middle row). Labelling of primary roots with MCB to visualize GSH, revealed low, but clearly increased GSH levels in *bir6 rml1* compared to *rml1* (**Fig. 7C** bottom row). Genotyping of dwarf seedlings in the *bir6 rml1* population after labelling of GSH and confocal imaging confirmed the expected double homozygosity. The combination of *rml1* with *gr1* had no effect on growth or GSH levels when compared to *rml1* single mutants (**Fig. 7C**).

**Fig. 7.**
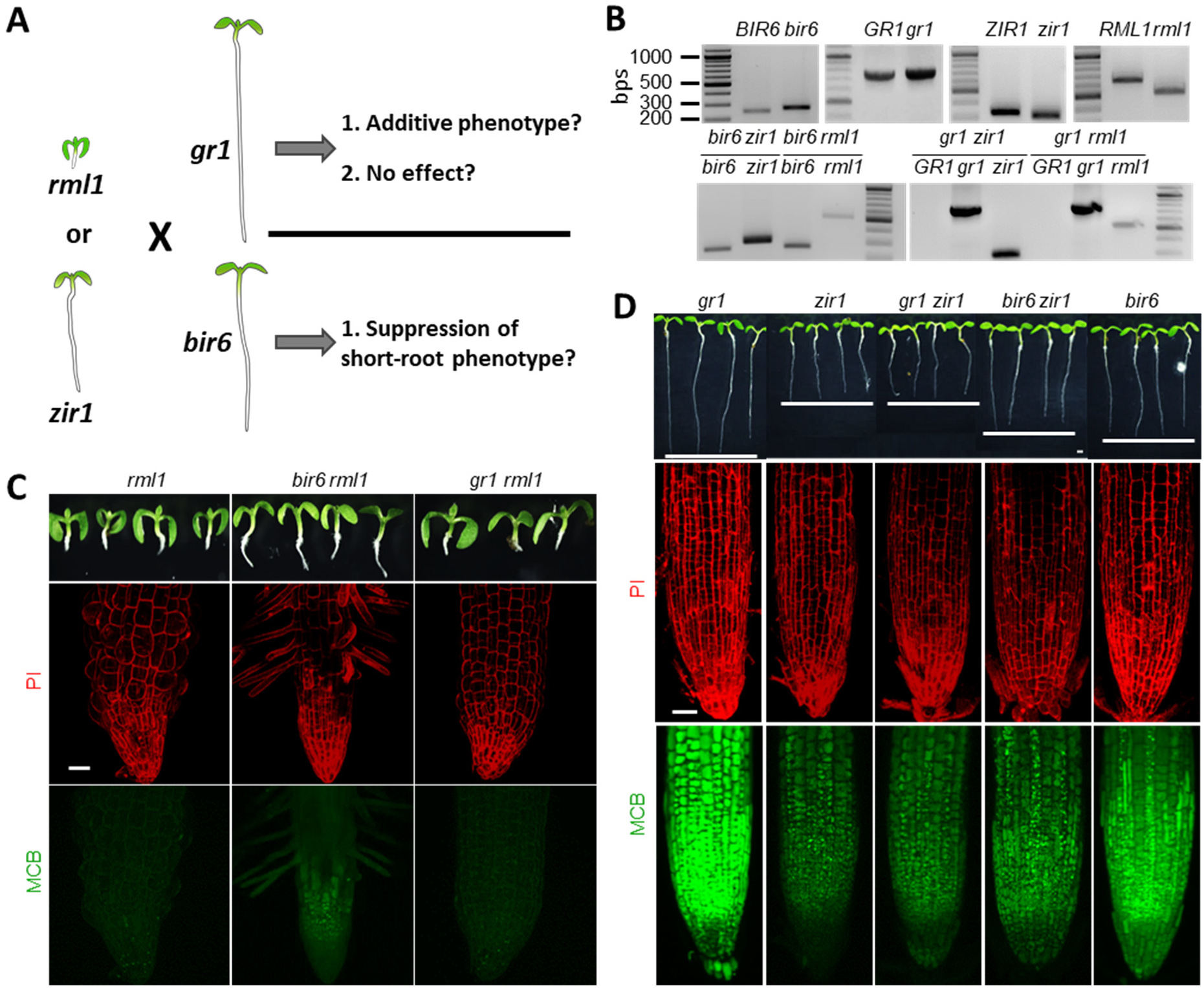
Suppression of *zir1* and *rml1* dwarf phenotypes by *bir6*. (A) Scheme of the genetic strategy used to test the hypotheses that root growth is controlled by the glutathione redox potential (*E*_GSH_) or by GSH levels. The severe dwarf mutants *rml1* and *zir1* were crossed with *gr1* and *bir6*, respectively. If *E*_GSH_ was the critical parameter for root growth, deletion of GR1 in low-GSH mutants should enhance the short root phenotype. Vice versa, if *E*_GSH_ was not critical, the combination with *gr1* should have little or no phenotypic effect. On the other hand, if the absolute GSH levels were critical, even a small increase in steady-state GSH levels, as introduced by crosses with *bir6*, should have a positive effect on root growth. (B) Isolation of double mutants. The upper panels show genotyping of heterozygous plants for the WT and mutant alleles. Restriction analysis for *bir6* (MfeI cuts the WT PCR fragment into 251 + 20 bps), *zir1* (HindIII cuts the mutant PCR fragment into 308 + 29 bps) and *rml1* (ApoI cuts the mutant PCR fragment into 540 + 127 bps) results in detectable fragment size differences. *gr1* is a T-DNA insertion mutant, for which the PCR fragments of the WT and the T-DNA allele are almost the same size. In the lower panels, PCR reactions confirmed the successful isolation of the respective double mutants in the double homozygous state. Note that no double bands were observed after the PCR fragments were restricted, confirming homozygosity for *zir1*, *rml1* and *bir6*. (C) Test for the effect of *bir6* and *gr1* on the phenotypic appearance of *rml1*. The upper panels show the phenotypes of seedlings five days after germination. The lower panels show the labelling of root tips with 100 µM MCB (green) and 50 µM PI (red) after incubation in dye solution for 30 min. (D) Test for the effect of *bir6* and *gr1* on the phenotypic appearance of *zir1*. The upper panels show the phenotypes of seedlings five days after germination. The lower panels show the labelling of root tips with 100 µM MCB and 50 µM PI after incubation in dye solution for 30 min. All fluorescence images are maximum projections of stacks of images collected on a confocal microscope. The GSB fluorescence obtained in *rml1*, *zir1* and the respective double mutants was generally very low. Therefore, the brightness of the images was artificially increased by 40%, to make the fluorescence visible. For *gr1* and *bir6* single mutants this was not necessary as the GSH level in these plants, and thus the GSB fluorescence intensity, were similar to the WT. Scale bars = 20 µm.

Unfortunately, the usability of *bir6 rml1* crosses was limited by the inability to maintain the mutant in double homozygous form. Therefore, we next turned to *zir1* and generated the corresponding crosses with *bir6* and *gr1*. *gr1* seedlings appear phenotypically like wild-type seedlings (Marty *et al*., 2009) and were hence used as a proxy for normal growth. In homozygous form, growth of *bir6* was slightly retarded (**Fig. 6A**; **Fig. 7D**), which is in agreement with previous observations (Koprivova *et al*., 2010a). The double homozygous *gr1 zir1* mutant was phenotypically indistinguishable from the *zir1* single mutant and the lack of GR1 activity did not impact on the amount of detectable GSH (**Fig. 7D**). In stark contrast, the *bir6 zir1* double mutant showed a clear suppression of the *zir1* dwarf phenotype, with primary roots reaching the length of *bir6* single mutant roots five days after germination. Reactivation of growth showed a clear correlation with increased GSH levels as shown by in vivo labelling with MCB (**Fig. 7D**). Taken together, the results obtained from these genetic experiments show that the GSH abundance rather than *E*_GSH_ is the limiting parameter for growth.

## Discussion

The tripeptide glutathione has several functions in plant metabolism, ranging from the detoxification of xenobiotics, ROS and heavy metals, to the post-translational modification of proteins. It was shown that the availability of glutathione is linked to root growth and cell-cycle control (Lee *et al*., 2025; Vernoux *et al*., 2000), but it remained unclear whether this is primarily associated with *E*_GSH_ or the absolute amount of glutathione. Here, we demonstrate that growth in both roots and shoots is controlled by the amount of GSH rather than the *E*_GSH_. Crosses between low-GSH mutants displaying different degrees of dwarfism and a mutant lacking cytosolic GR1, showing a WT-like phenotype, did not enhance the growth-defective phenotypes. In contrast, an increase in GSH amounts resulting from crosses between low-GSH mutants and a mutant with a slightly increased steady-state GSH resulted in attenuation of the phenotype.

### Partial glutathione deficiency causes impaired growth of shoots and roots

We initially investigated the cause of diminished growth observed in severely glutathione-deficient mutants by using a collection of six allelic *gsh1* mutants, which all carry mutations in close proximity to the GSH1 active site that lead to different degrees of glutathione deficiency. Consistent with the respective original reports, seedlings of *rax1*, *pad2*, *cad2*, and *nrc1*, did not show any obvious impairment in primary root growth five days after germination. A pronounced short-root phenotype was observed only in *zir1*, and post-embryonic meristem activity in the *rml1* roots was almost completely abolished (**Fig. 1**). However, roots analysed for their morphological parameters 20 days after germination showed a small decrease in total root length even in seedlings with intermediate GSH levels. A similar observation was reported earlier for primary roots of seedlings 10 days after germination (Bashandy *et al*., 2010; Marquez-Garcia *et al*., 2014; Urbancsok *et al*., 2018). This decrease in total root length is primarily due to a reduction in lateral root length (**Fig. 2**). In contrast to earlier reports by Bashandy *et al*. (2010) and Marquez-Garcia *et al*. (2014), the number of lateral roots was not significantly affected in *rax1*, *pad2* and *cad2*, and only to a minor degree in *nrc1.* This suggests that the formation of lateral root primordia and their further development is not generally compromised in plants with intermediate GSH levels. This was also apparent when the severely glutathione-deficient mutants *zir1* and *rml1* were rescued with external supply of 1 mM GSH. In this case, lateral root primordia were clearly visible on the primary root of *rml1*, but did not yet form proper laterals within four days on GSH (**Fig. 4C**). The initiation of lateral roots requires the development of local auxin maxima in the lateral root primordia (Du and Scheres, 2018). The dependence of primordia formation on GSH is consistent with earlier observations that inhibiting GSH biosynthesis with BSO represses the expression of multiple auxin transporters and auxin response genes (Bashandy *et al*., 2010; Koprivova *et al*., 2010b).

At rosette stage, all mutants showed a smaller shoot than the wild-type control. This suggests that even partial GSH deficiency in *cad2*, *pad2*, *rax1* and *nrc1* affect vegetative growth performance. Whether this growth restriction results from direct effects of glutathione on the shoot meristem deserves further investigation. Yet, it would be highly surprising if a regulatory effect of glutathione on cell cycle activity was restricted to the root meristem.

### Severe glutathione deficiency causes a global increase in amino acids

The amount of glutathione in Arabidopsis is inversely correlated with the amount of cysteine, with cysteine levels reaching approximately seven times higher values in *rml1* compared to the WT (**Fig. 3C, D**), which is slightly higher than a 5-fold increase reported for *GSH1* antisense lines (Xiang *et al*., 2001). Such free cysteine is potentially toxic as it may cause iron reduction which, in the presence of H_2_O_2_, would set off a Fenton reaction and lead to the formation of highly toxic hydroxyl radicals (Meyer and Hell, 2005). Production of these radicals as a consequence of elevated cysteine levels in *rml1* could cause growth inhibition through diverse inhibitory effects on proteins and DNA. Such effects would be expected to be accompanied by severe membrane damage, but the absence of nuclear DNA-staining with PI in *rml1* roots thus speaks against a link between the observed growth defects and elevated cysteine levels. Similarly, the ability to reactivate cell division and growth of *rml1* by feeding GSH argues against DNA damage that otherwise would frequently lead to cell death (Serrano-Mislata *et al*., 2025). This is further supported by the observation that feeding Arabidopsis seedlings with up to 1 mM cysteine has no apparent phenotypical effect on root growth (Ravelo-Ortega *et al*., 2021) The same authors also tested all the other proteinogenic amino acids for their ability to repress growth and found that 11 amino acids affected the growth of primary roots by at least 40%, with lysine and tryptophan being the most effective inhibitors. Amino acid profiling in *rml1* revealed a roughly twofold increase in the levels of all recorded amino acids (**Fig. 5**), including those reported to cause growth repression to some degree (Ravelo-Ortega *et al*., 2021). It is possible that the additive effects of different amino acids could ultimately cause complete growth inhibition in *rml1* roots. However, similar measurements in *zir1* revealed a wild-type amino acid profile (see **Fig. 5**), suggesting that the partial growth repression in *zir1* and the complete inhibition in *rml1* are most likely independent of the accumulating amino acids. The pronounced accumulation of all amino acids in *rml1* is more likely caused by a block in protein biosynthesis resulting in less demand for amino acids as well as proteolysis and autophagy (Hildebrandt, 2018). Thus, the results support the conclusion that the evident growth repression observed in all mutants of the *GSH1* allelic series appears to be directly linked to glutathione.

### An oxidative shift in cytosolic *E*_GSH_ does not affect primary root growth

To further probe the function of glutathione in growth, we considered both, the *E*_GSH_ and the absolute amount of GSH. Unfortunately, it is not possible to disentangle the two parameters, because the glutathione redox pair comprises two GSH molecules which form one GSSG molecule when two electrons are removed. Due to this 2:1 stoichiometry, changes in both the redox state and the absolute amount of glutathione affect *E*_GSH_. In subcellular compartments containing GRs, the local *E*_GSH_ is highly negative because strong GR activity leaves only low nanomolar concentrations of GSSG in a background of low millimolar concentrations of GSH, provided that sufficient NADPH is available as a reducing agent (Marty *et al*., 2019; Marty *et al*., 2009; Meyer *et al*., 2007; Schwarzländer *et al*., 2008). Assuming 2.5 mM total glutathione in the cytosol of wild-type plants and a degree of oxidation of 0.002% for the glutathione pool, *E_G_*_SH_ would be approximately -310 mV (Meyer *et al*., 2007). For *cad2* grown under the same growth conditions, 25% of the wild-type glutathione (**Fig. 3C**) falls well into the range of 15-30% glutathione reported earlier (Howden *et al*., 1995). Assuming the same degree of oxidation of 0.002% set by GR1 and sufficient NADPH supply, the Nernst equation would predict an *E*_GSH_ of -293 mV, which matches well with the fluorescence ratio obtained for cytosolic roGFP2 in the root tip (**Fig. 3E**) and confirms earlier measurements(Meyer *et al*., 2007). In the other mutants with intermediate glutathione levels, roGFP2 is oxidised to a similar degree (**Fig. 3E**). Differences in the glutathione level as measured by HPLC, e.g., between *rax1* and *cad2*, cannot be easily resolved with roGFP2 (**Fig. 3C, E**) because a difference between 20 and 30% of wild-type glutathione levels would cause only a 5 mV shift in *E*_GSH_. It should also be noted here that HPLC measurements were taken from whole seedlings, whereas roGFP2 measurements were taken exclusively from root tips. Therefore, comparisons of *E*_GSH_ values calculated based on glutathione amounts measured by HPLC and *E*_GSH_ values derived from roGFP2 measurements can only ever be approximate. Assuming 375 µM glutathione and a maintained degree of oxidation of 0.002% in *zir1*, this would result in an *E*_GSH_ of approximately - 286 mV. Similarly, an assumption of 125 µM glutathione in *rml1* would result in an *E*_GSH_ of -272 mV. However, the roGFP2 fluorescence ratios measured in these two mutants indicate even less negative values. This is consistent with earlier measurements in *rml1*, which, based on the use of two probe variants, suggested a consensus *E*_GSH_ of -260 mV in the cytosol (Aller *et al*., 2013). These deviations between measurements and theoretical calculations might be explained by metabolic deficiencies and a compromised supply of NADPH in these mutants. For *rml1*, at least, a metabolic problem is evident from the amino acid profile and the complete growth arrest. Similarly, the decreased growth in *zir1* might also be related to decreased metabolic flux even if steady state levels of amino acids in this case could still be maintained. Whether such metabolic problems are cause or consequence remains unknown at this point.

The deletion of GR1 in the cytosol causes a shift of *E*_GSH_ towards less negative values of at least 20 mV, with no significant change in the total amount of glutathione or the amount of GSH (Marty *et al*., 2009). The deletion of GR1 in *zir1* and *rml1* would necessarily cause *E*_GSH_ to shift further towards less negative values. For *zir1 gr1*, this would mean that *E*_GSH_ should decrease to values approaching those found in the *rml1* cytosol. The absence of any additive phenotype in *zir1 gr1* and *rml1 gr1* double mutants compared to the *zir1* and *rml1* single mutants, indicates that growth is not restricted or controlled by *E*_GSH_ in the cytosol. A direct repressive effect of a low *E*_GSH_ on the cell cycle is also unlikely, because cell division in embryos proceeds with very low amounts of GSH. Labelling of U-turn stage embryos with MCB indicated that homozygous *gsh1* null mutant embryos contain only minute amounts of GSH, which is supplied by maternal tissues (Cairns *et al*., 2006; Lim *et al*., 2011). Although not measured directly, the *E*_GSH_ in these embryos can be expected to be even less negative than in *rml1* seedlings, arguing against control of the cell cycle by *E*_GSH_ or an *E*_GSH_-related process.

### Primary root growth is dependent on the absolute amount of glutathione

Compared to the effect of *GR1* deletion, small changes in the steady state level of glutathione have only a minor impact on the cytosolic *E*_GSH_. The steady state level of glutathione is defined by continuous biosynthesis on one hand and consumption, particularly for detoxification of H_2_O_2_ and xenobiotics, on the other hand (Noctor *et al*., 2011; Ugalde *et al*., 2025). In Brassicaceae, GSH is also consumed for the biosynthesis of sulfur-containing secondary metabolites like glucosinolates and camalexin (Geu-Flores *et al*., 2011). In addition, continuous degradation of GSH by γ-glutamyl peptidases (GGPs) has been reported and even though direct degradation appears to create a futile cycle of ATP-consuming biosynthesis and degradation, deletion of GGP1 results in an increase of total glutathione by 30% (Ito *et al*., 2022). With the goal of inducing only minor shifts in total glutathione, we selected *bir6*, which maintains a slightly higher glutathione pool after treatment with the biosynthetic inhibitor BSO (**Fig. 6**). Even though the cause for the decreased consumption of GSH in *bir6* is not understood (Koprivova *et al*., 2010a), the mutant allowed to test the hypothesis for the role of the absolute glutathione amount on cell cycle activity and growth. Partial de-repression of primary root growth in *rml1 bir6* and *zir1 bir6* is correlated with a slight increase of GSH in root tips compared to the respective single mutants (**Fig. 7**). In *rml1*, this de-repression is visible by slightly longer primary roots that indicate reestablishment of cell cycle activity, albeit at very low level. The more pronounced effect is on the growth of root hairs, which supports a role of GSH in root hair growth (Sánchez-Fernández *et al*., 1997; Trujillo-Hernandez *et al*., 2020).

## Conclusion

Taken together, our data demonstrate that the absolute level of glutathione rather than *E*_GSH_ governs the meristematic activity in the primary root of Arabidopsis. Based on the partial suppression of short-root phenotypes of GSH-deficient *zir1* mutants by combination with *bir6*, we conclude that a small increases in the steady state levels of GSH is sufficient to either minimize deleterious effects from endogenous metabolites or to activate yet unknown components in the cell cycle. The absence of any obvious phenotypic effect of *gr1* on the GSH-deficient mutants *zir1* and *rml1*, also suggests that ROS, and in particular H_2_O_2_, the detoxification of which through the ascorbate-glutathione pathway may lead to a transient increase in GSSG, are unlikely to play a key role in cell cycle control; at least not mediated via glutathione. Our finding that maintenance of GSH amounts as decisive factor provides an instructive foundation for further research to focus on the multiple functions of glutathione in detoxification of endogenous toxic metabolites. Candidate pathways that may require GSH amounts above certain thresholds are the detoxification pathway for methylglyoxal, which involves GSH as a cofactor (Cassier-Chauvat *et al*., 2023), iron-sulfur cluster transfer by glutaredoxins, which relies on GSH as a cofactor (Moseler *et al*., 2015), or the detoxification of nitric oxide (NO), which involves a non-catalysed second order attack of GSH on NO to form GSNO for further reduction (Groß *et al*., 2013).

## Author contributions

AJM, MS, SKo, JPR: conceptualization; SAKB, JMU, KN, MJ, SKr, AM, MTS: investigation; SAKB, MTS, JMU, SW, AM: formal analysis; AJM: writing—original draft preparation; AJM, SW, SKo, JPR: writing— review and editing; AJM: funding acquisition; AJM, MS: supervision.

## Acknowledgements

We thank Bastian Welter (University of Cologne) for technical assistance with HPLC measurements, Bernd Kastenholz, Silvia Braun and Thorsten Brehm (Forschungszentrum Jülich) for performing the root and shoot phenotyping experiments at Forschungszentrum Jülich.

## Supplementary data

Table S1. Oligonucleotides and restriction analysis used for genotyping.

Table S2. Amino acids in wildtype, *zir1* and *rml1*.

---

**Supplemental Table S1.** Oligonucleotides and restriction analysis used for genotyping.

| Allele | Type of mutation | Forward primer | Reverse primer | Fragment size (bp) | Restriction enzyme | Fragment A | Fragment B |
| --- | --- | --- | --- | --- | --- | --- | --- |
| *GR1* | Wild type | *cs330* | *cs331* | 763 |  |  |  |
| *gr1-1* | T-DNA | SALK_LBb1.3 | *cs331* | ~800 |  |  |  |
| *zir1* | Point mutation | *cs3026* | *cs3027* | 337 | HindIII | 308 | 29 |
| *rml1* | Point mutation | *cs641* | *cs642* | 667 | ApoI | 540 | 127 |
| *bir6-1* | Point mutation | *cs2985* | *cs2986* | 271 | MfeI/MunI | 251 | 20 |

**Supplemental Table S2.**
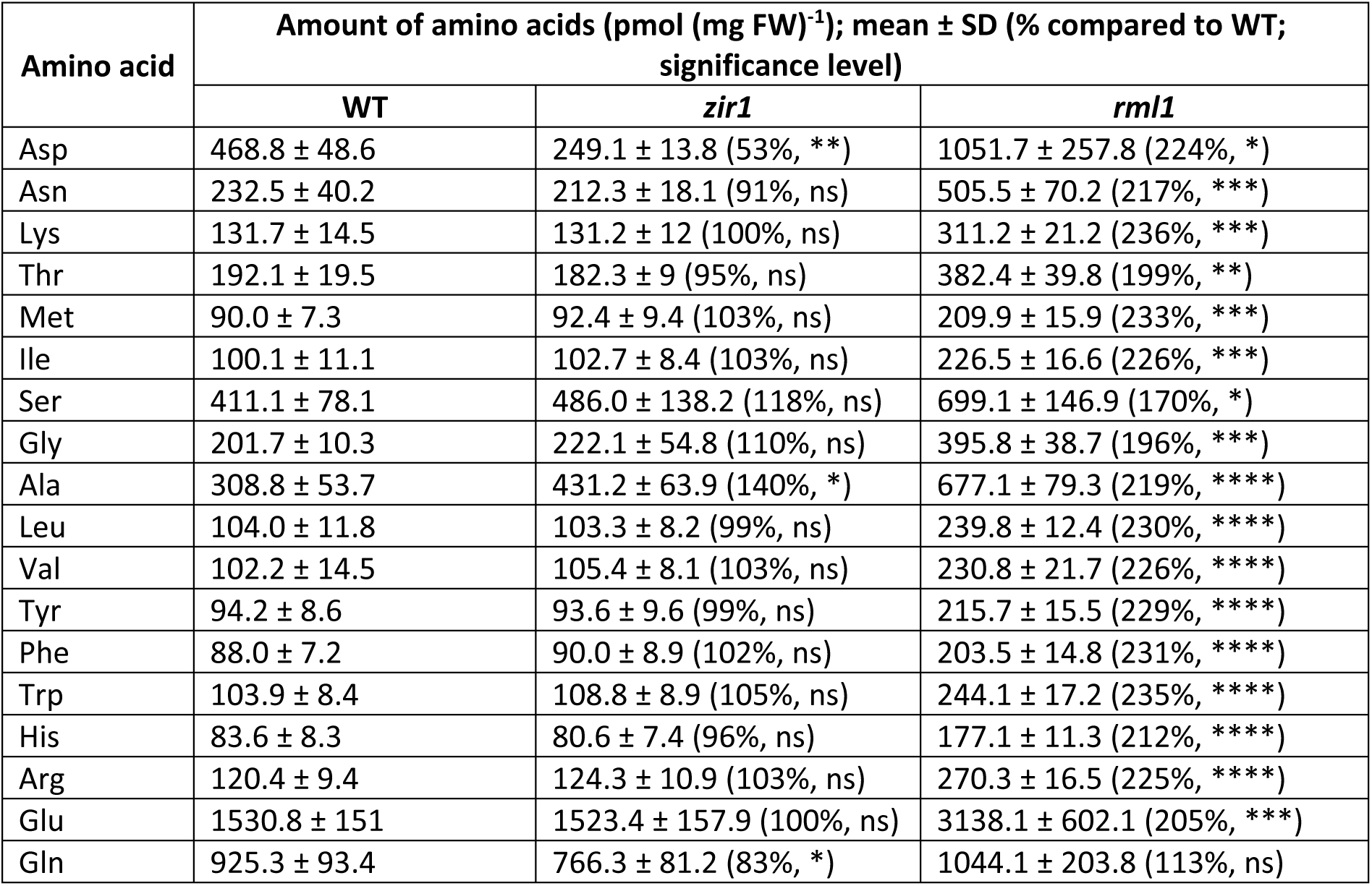
Content of amino acids of Arabidopsis WT, *zir1*, and *rml1*. The statistical analysis (one-way ANOVA with Dunnett’s multiple comparisons test for WT vs. *gsh1* mutant) indicated significance levels; *n* = 4-5; *P ≤ 0.1, **P ≤ 0.01; ***P ≤ 0.001; ****P ≤ 0.0001, ns: not significant.

| Amino acid | Amount of amino acids (pmol (mg FW)<sup>-1</sup>); mean ± SD (% compared to WT; significance level) |  |  |
| --- | --- | --- | --- |
|  | WT | *zir1* | *rml1* |
| Asp | 468.8 ± 48.6 | 249.1 ± 13.8 (53%, **) | 1051.7 ± 257.8 (224%, *) |
| Asn | 232.5 ± 40.2 | 212.3 ± 18.1 (91%, ns) | 505.5 ± 70.2 (217%, ***) |
| Lys | 131.7 ± 14.5 | 131.2 ± 12 (100%, ns) | 311.2 ± 21.2 (236%, ***) |
| Thr | 192.1 ± 19.5 | 182.3 ± 9 (95%, ns) | 382.4 ± 39.8 (199%, **) |
| Met | 90.0 ± 7.3 | 92.4 ± 9.4 (103%, ns) | 209.9 ± 15.9 (233%, ***) |
| Ile | 100.1 ± 11.1 | 102.7 ± 8.4 (103%, ns) | 226.5 ± 16.6 (226%, ***) |
| Ser | 411.1 ± 78.1 | 486.0 ± 138.2 (118%, ns) | 699.1 ± 146.9 (170%, *) |
| Gly | 201.7 ± 10.3 | 222.1 ± 54.8 (110%, ns) | 395.8 ± 38.7 (196%, ***) |
| Ala | 308.8 ± 53.7 | 431.2 ± 63.9 (140%, *) | 677.1 ± 79.3 (219%, ****) |
| Leu | 104.0 ± 11.8 | 103.3 ± 8.2 (99%, ns) | 239.8 ± 12.4 (230%, ****) |
| Val | 102.2 ± 14.5 | 105.4 ± 8.1 (103%, ns) | 230.8 ± 21.7 (226%, ****) |
| Tyr | 94.2 ± 8.6 | 93.6 ± 9.6 (99%, ns) | 215.7 ± 15.5 (229%, ****) |
| Phe | 88.0 ± 7.2 | 90.0 ± 8.9 (102%, ns) | 203.5 ± 14.8 (231%, ****) |
| Trp | 103.9 ± 8.4 | 108.8 ± 8.9 (105%, ns) | 244.1 ± 17.2 (235%, ****) |
| His | 83.6 ± 8.3 | 80.6 ± 7.4 (96%, ns) | 177.1 ± 11.3 (212%, ****) |
| Arg | 120.4 ± 9.4 | 124.3 ± 10.9 (103%, ns) | 270.3 ± 16.5 (225%, ****) |
| Glu | 1530.8 ± 151 | 1523.4 ± 157.9 (100%, ns) | 3138.1 ± 602.1 (205%, ***) |
| Gln | 925.3 ± 93.4 | 766.3 ± 81.2 (83%, *) | 1044.1 ± 203.8 (113%, ns) |

